# Habituation and individual variation in the endocrine stress response in the Trinidadian guppy (*Poecilia reticulata*)

**DOI:** 10.1101/337006

**Authors:** T.M. Houslay, R.L. Earley, A.J. Young, A.J. Wilson

**Affiliations:** Centre for Ecology and Conservation, University of Exeter (Penryn Campus), Cornwall, TR10 9FE, UK; Department of Biological Sciences, University of Alabama, Biology Building 211-213, Box 870344, Tuscaloosa, AL 35487, USA

## Abstract

The vertebrate stress response enables individuals to react to and cope with environmental challenges. A crucial aspect of the stress response is the elevation of circulating glucocorticoids. However, continued activation of the stress response under repeated (or chronic) stress can be damaging to fitness. Under certain circumstances it may therefore be adaptive to habituate to repeated exposures to a particular stressor by reducing the magnitude of any associated release of glucocorticoids. Here, we investigate whether Trinidadian guppies (*Poecilia reticulata*) habituate to repeated exposure to a mild stressor, using a waterborne hormone sampling approach that has previously been shown to elicit a stress response in small fish. We also test for individual variation in the extent of habituation to this stressor. Concentrating on freely circulating cortisol, we found that the first exposure to the assay induced high cortisol release rates but that guppies tended to habituate quickly to subsequent exposures. There were consistent differences among individuals in their average cortisol release rate (after accounting for effects of variables such as body size) over repeated exposures. Our analyses did not find evidence of individual differences in habituation rate, although limitations in statistical power could account for this finding. We also present data on free 11-ketotestosterone, in addition to conjugated forms of both hormones. We discuss consistent individual differences around the general pattern of habituation in the flexible stress response, and highlight the potential for individual variation in habituation to facilitate selection against the deleterious effects of chronic stress.

**Summary statement:** Trinidadian guppies habituate quickly to repeated stress exposure, and exhibit consistent differences in their endocrine stress response. We provide a framework for analysing individual variation in habituation rate.

## Introduction

Stress responses involve a complex suite of behavioural, neuroendocrine, and physiological processes that act to maintain organismal health and homeostasis in the face of unpredictable environmental challenges (Selye 1973; Korte *et al*. 2005; Romero, Dickens & Cyr 2009). While disagreements over terminology persist (McEwen and Wingfield, 2010; Romero et al., 2009), stress response mechanisms are broadly seen as underlying a process of “achieving stability through change” (McEwen and Wingfield, 2003) and are vital for dealing with acute environmental challenges. However, these same mechanisms can actually be damaging to fitness when organisms are exposed to chronic stressors (Boonstra, 2013; Dallman et al., 1992; Huether, 1996). Given the expectation of such causal links between stress response and fitness, there has been increasing interest in characterising variation among individuals for both behavioural and neuroendocrine stress response phenotypes (e.g., Cockrem 2013; Houslay *et al*. 2017). Variation among individuals is a prerequisite for selection to occur, and, if such variation also includes a genetic component, then stress response traits have the potential to evolve under natural selection in wild populations (Atwell *et al.* 2012; Jenkins *et al*. 2014) or be targeted by artificial selection strategies in captive ones (Pottinger & Carrick 1999; Louison *et al.* 2017).

Among-individual variation has now been documented in behavioural (Bell et al., 2009) and neuroendocrine (Hau et al., 2016) traits across many taxa. For neuroendocrine traits in particular, a distinction is commonly made between ‘baseline’ and ‘reactivity’ phenotypes. The former represents the phenotypic state (e.g. circulating glucocorticoid (GC) level) of an ‘unstressed’ individual, while the latter is the plastic change in expression (e.g. increase in GC level) when challenged by an acute stressor. Both aspects are likely to vary among individuals within a population (Williams, 2008), and their recognition as distinct-but potentially correlated-traits allows a more nuanced view of how selection might shape the stress response (Bonier and Martin, 2016; Cox et al., 2016; Hau and Goymann, 2015; Romero, 2004; Taff and Vitousek, 2016). Nonetheless, we suggest this framework will not be sufficient to understand the fitness consequences of chronic stress exposure. This is because plasticity is relevant not just to ‘reactivity’ when challenged by an acute stressor, but also to longer-term changes in phenotypic state when faced by chronic repeated exposures to stressors. For instance, stress-induced GC release can result in the phenomenon of sensitisation, where the individual enters a state of hyper-responsiveness to the same or novel stimuli (Belda et al., 2015). A more common response to repeated exposure to the same acute stressor, however, is habituation (Martí and Armario, 1998).

Habituation can be defined as occurring where repeated, or chronic, application of a stimulus results in a progressively weaker stress response (Thompson and Spencer, 1966). Intuitively, habituation to chronic stressors may offer a route to improved health, particularly if simple behavioural responses (e.g. avoidance of the stressor) are curtailed, which will commonly be the case in captive populations. For example, under chronic stress exposure, continued activation of the hypothalamic-pituitary-adrenal/interrenal (HPA/I) axis can occur. This is a metabolically costly response system (Korte et al., 2005) and sustained elevation of circulating glucocorticoids (GCs) is known to reduce growth, suppress immune function and reproduction, and increase mortality (Berga, 2008; Hodgson et al., 2012; Segerstrom and Miller, 2004; Young et al., 2006). As such it should be adaptive for an organism to habituate (via reductions in the magnitude of the HPA/I response) to stressors that are not inherently harmful (Grissom and Bhatnagar, 2009; McEwen, 2001). If present, variation among individuals in their ability to habituate to a repeated stressor thus represents another form of variable plasticity that could facilitate selection to reduce the deleterious effects of chronic stress in natural and captive populations.

In this study we seek to characterise among-individual variation in stress-related endocrine state and habituation in the Trinidadian guppy, *Poecilia reticulata*. We focus principally on HPI activity and thus cortisol, the major GC in teleost fish (Mommsen et al., 1999). We also conduct a parallel analysis of 11-ketotestosterone (subsequently 11KT) from the samples collected. Links between 11KT and stress response have previously been reported in fishes; chronically elevated cortisol can inhibit 11KT synthesis (Consten et al., 2001; Consten et al., 2002), and social/isolation stressors have been shown to reduce 11KT while elevating cortisol (Galhardo and Oliveira, 2014; Haddy and Pankhurst, 1999; Kubokawa et al., 1999). Here we separate both of these target hormones into their ‘free’ and ‘conjugated’ fractions for analysis. In the main text we present only results from free hormones, the concentration of which in the water is taken to scale with the ‘physiologically active’ concentration of hormones in the fish’s circulation across the duration of the sampling period (Scott and Ellis, 2007). We refer interested readers to Appendix A for information on the conjugated fractions.

Our objectives are not only to determine whether individual fish differ consistently in stress-related endocrine state, but also to determine whether habituation occurs and, if so, whether rates vary among individuals. Our focus on among-individual differences requires repeated sampling of individuals; since this is not feasible using blood sampling in guppies, we adopt a less invasive, waterborne approach to characterising endocrine state. While this method has been widely validated and used for studies of small fishes (reviewed in Scott & Ellis 2007; Scott *et al*. 2008), including guppies (Fischer et al., 2014), the handling required to transfer the fish into the confined space in which waterborne sampling will be conducted causes cortisol release, complicating attempts to measure baseline GC (e.g., Wong et al., 2008). However, in the current context, this is actually advantageous as we use this initial handling and confinement itself as the stressor to which habituation is predicted to occur. For free cortisol we predict that: (i) average levels will decline over repeated handling and sampling events (1 every 48 hours, for a total of 4 repeats), consistent with habituation; (ii) there will be variation among individuals around the average response, with individuals differing in both average ‘reactive’ endocrine state and their rate of habituation with repeated subsequent exposures; and (iii) that individuals with higher average circulating cortisol levels also habituate at a slower rate. We make no specific predictions about free 11KT levels beyond a general expectation that they will be negatively correlated with free cortisol.

## Methods

### Animal husbandry and welfare

The guppies used in this study were taken from our captive population, descended from wild fish collected in 2008 from the lower Aripo River, Trinidad, and housed at the University of Exeter’s Penryn campus. We sampled 32 adult fish (17 females, 15 males) from a stock tank haphazardly, and tagged each for identification purposes with coloured elastomer injection under mild MS222 sedation. We housed fish in 2 mixed-sex groups of equal size, using separate tanks that shared a common recirculating water supply. All fish were fed to satiation twice daily (0800-1000h and again at 1600-1800h) using commercial flake food and live *Artemia nauplii*. The experiment described here was carried out in accordance with the UK Animals (Scientific Procedures) Act 1986 under licence from the Home Office (UK), and with local ethical approval from the University of Exeter.

### Sampling protocol & hormone assay

We used a waterborne sampling method (Scott *et al*. 2008) to obtain 4 repeated measures of hormones from individual fish at 48hr intervals. To control for diel fluctuations in cortisol, we collected all samples between 1200-1400h. In all cases, we first netted an entire group from the housing aquarium quickly using a large net and transferred them to 2 holding tanks for moving to an adjacent quiet room (performed within 20 seconds of the net first hitting the water). We then transferred fish to individual beakers containing 300ml of clean water taken from the main supply (which serves as input to the main housing units) for sample collection, placing beakers within cardboard ‘chambers’ to prevent fish from seeing each other or being disturbed. 1 fish was transferred every 30 seconds, alternating across holding tanks, such that all fish were in their beakers within 10 minutes of the initial netting (with maximum 11 net entries into the water). Individual ID was noted during transfer to the beaker, and sampling order retained for use in statistical models to control for effects of both the time between first net entry and transfer to the beaker and the number of net entries into the water. After 60 minutes in the beaker, we removed each individual by pouring its sample through a clean net into a second beaker (and returned fish to their home tank immediately). At each time point, we also collected 2 ‘blank’ samples from the water supply.

Immediately after sampling, we filtered the water samples (Grade 1 filter paper, Whatman) and then passed them through solid phase C18 extraction columns (Sep-Pak C18 3cc, Waters) that had been primed with 2 × 2ml HPLC-grade methanol followed by 2 × 2ml distilled water, keeping the column moist prior to sample extraction. We used high-purity tubing (Tygon 2475, Saint-Gobain) to pass the water samples to the columns, through which they were drawn slowly under vacuum pressure (see Earley et al., 2006 for detail). Following extraction, we washed columns with 2 × 2ml distilled water to purge salts. We then covered both ends of each column with film (Parafilm M, Bemis) and stored them at −20°C for future analysis. We rinsed all beakers, tubes and funnels with 99% ethanol and distilled water prior to each sampling procedure.

We later shipped the frozen columns on dry ice from the University of Exeter’s Penryn campus to the University of Alabama. We thawed each column and purged with 2 x 2ml distilled water before eluting both free and conjugated (sulphated and glucuronidated) hormones. We used 2 x 2ml ethyl acetate and 2 × 2 ml HPLC-grade methanol in successive, separate washes to elute free and conjugated hormones respectively into separate 13 × 100 mm borosilicate vials. For the free fraction, ethyl acetate was evaporated under nitrogen in a water bath at 37°C, leaving a hormone residue, which was resuspended in 600 μl of 5% EtOH: 95% EIA kit buffer (the latter supplied with Cayman Chemicals, Inc. EIA kits) and vortexed for 20 minutes. For the conjugated hormones, methanol was evaporated under nitrogen and the hormone residue was resuspended in 600 μl of 5% EtOH: 95% EIA kit buffer. The resuspended conjugates were then diluted with 8 ml distilled water, re-extracted over primed solid phase C18 extraction columns (Sep-Pak C18 3cc, Waters) and eluted into 13 x 100 mm borosilicate vials with 2 x 2 ml successive washes of methanol, which was evaporated under nitrogen. We processed these samples further according to previously published protocols with some modification (Canario and Scott, 1989; Scott and Canario, 1992; Scott and Vermeirssen, 1994).

Briefly, 1 ml trifluoroacetic acid [TFA]:ethyl acetate (1:100) was added to the hormone residue and incubated overnight in a water bath at 37°C. The TFA:ethyl acetate was evaporated under nitrogen at 37°C followed by addition of 0.5 ml 0.1M sodium acetate buffer (pH 4.5). We vortexed the sample for 5 minutes, followed by addition of 4 ml diethyl ether and a further 5 minutes of vortexing. We then allowed samples to sit for approximately 30 minutes to enable separation of the organic and aqueous phases, and then immersed the samples in a methanol-dry ice bath to freeze the aqueous phase. The organic phase containing the sulphated fraction was poured off into a new 13 × 100 mm borosilicate vial. 10μl of β-glucuronidase (Sigma Aldrich, Cat No. G-7017) was added to the remaining aqueous phase and incubated overnight in a water bath at 37°C, followed by the diethyl ether extraction as described above. The organic phase containing the glucuronidated fraction was poured into the same vial as the ether containing the sulphated fraction to produce a sample with total conjugated hormone. Diethyl ether was evaporated under nitrogen in a 37°C water bath and the hormone residue was resupended in 200μl of 5% EtOH: 95% EIA kit buffer followed by vortexing for 20 min. We ran assays in strict accordance with the manufacturer’s instructions, and all samples were run in duplicate.

We generated a pooled waterborne hormone extract for males and females separately by combining 20 μl taken from each of the 128 samples. These pools were serially diluted to assess parallelism with the standard curves from the Cayman Chemicals, Inc cortisol and 11KT kits. Two serial dilution curves were run in duplicate for each hormone, and each was parallel to the standard curve (slope comparison test, Zar, 1996, p. 355; Cortisol-1: t_12_=0.204, *P* = 0.84; Cortisol-2: t_12_=0.013, *P* = 0.99; 11KT-1: t_12_=0.02, *P* = 0.98; 11KT-2: t_12_=0.005, *P* = 0.99). We also ran the pooled samples in duplicate at both the beginning and end of each of the four 96-well plates for each hormone (free fraction) to assess intra-and inter-assay coefficients of variation. Intra-assay coefficients of variation were 6.9%, 11.5%, 4.9%, and 9.5% for each of the cortisol plates and 7.8%, 3.4%, 2.9%, and 1.6% for each of the 11KT plates; the inter-assay coefficients for cortisol and 11KT were 10.7% and 7.3%, respectively. We generated a separate pool for the conjugated hormone, which was run in duplicate at the beginning and end of an additional four 96-well plates for each hormone (conjugated fraction). Intra-assay coefficients of variation were 4.9%, 7.7%, 6.6% and 5.6% for cortisol and 5.6%, 15.2%, 7.6% and 10.1% for 11KT; the inter-assay coefficients for cortisol and 11KT were 6.5% and 11.3%, respectively.

### Modelling approach and statistical analysis

In what follows we conceptualise habituation as plasticity in endocrine state with respect to repeated stressor exposure. This allows us to test for and characterise among-individual variation in habituation using the well-known ‘character state’ and ‘reaction norm’ frameworks for analysing plasticity (Henderson, 1982; Nussey et al., 2007; Van Noordwijk, 1989; Via et al., 1995). In the reaction norm framework, which we illustrate graphically in Fig. 1, each individual’s endocrine state is modelled as a linear function of sampling repeat number. This allows us to characterise among-individual variation in habituation (i.e. plasticity in endocrine state) using random regression (i.e. random slope) linear mixed effect models. Under the alternative character state approach, endocrine state at each sampling repeat is treated as a distinct (but potentially correlated) sub-trait. When more than 2 such sub-traits exist, the character state approach provides more information than the simple linear formulation of random regression, although this additional flexibility does require the estimation of additional parameters (Roff and Wilson, 2014). Using this approach, positive (within-individual) correlation across observations provides evidence of repeatable variation in endocrine state. If variation in habituation also occurs then this will lead to changes in observed endocrine state variance over sampling repeats, and/or declining (within-individual) correlation with increasing time between observations (such that *r*_1,2_ > *r*_1,3_ > *r*_1,4_).

**Figure 1:**
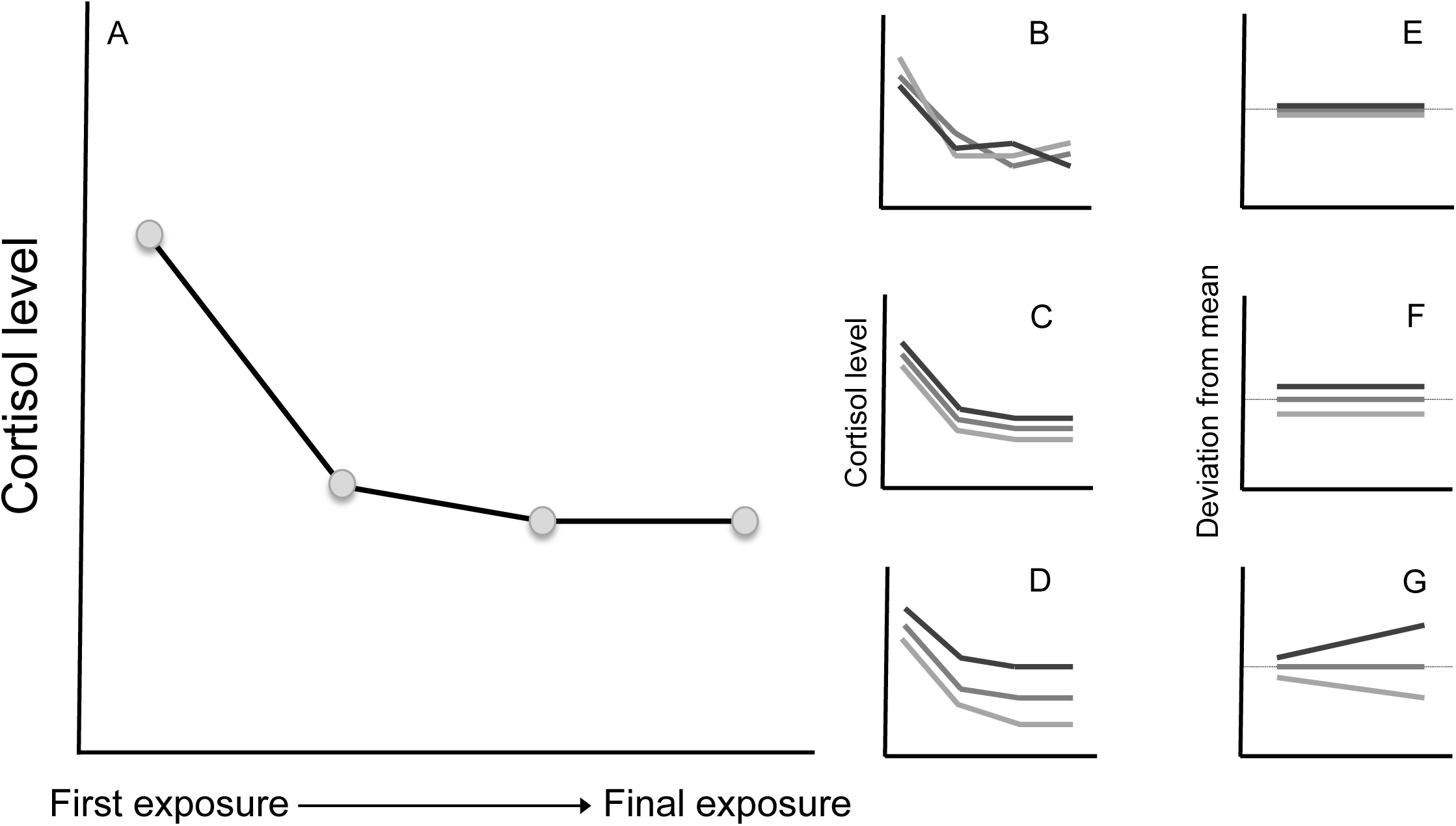
Characterising individual variation in habituation. Main panel (A) shows how habituation across repeated stressor exposures is predicted to affect average levels of cortisol (adapted from Fig. 3 of Fischer et al., 2014). Inset panels show examples of three individuals sampled haphazardly from such a population, indicating (B) no consistent differences among individuals, (C) among-individual differences in intercept only (average cortisol levels differ but rate of habituation does not), (D) among-individual differences in both intercept and slope (variation in both average cortisol levels and the rate of habituation). Note that while absolute cortisol levels depicted here do not follow linear (i.e., straight line) reaction norms, this assumption is not made in our analysis. Rather we assume only that a straight line is adequate to describe each individual’s deviation from the replicate-specific mean, as is the case in this depiction (E-G; dotted black line indicates zero deviation from the population mean).

The analysis procedure described below was performed separately for both free and conjugated forms of cortisol (log nanograms) and 11-ketotestosterone (log picograms). We present only the results for free hormones below, with results and discussion of conjugated forms available in Appendix A. All analyses were performed in R version 3.4.1 (R Core Team, 2017), using the package ASreml-R 3.0 (Butler, 2009) to test for hypothesised changes in mean endocrine state with repeated samplings, and to characterise variation among-individuals around this mean state. We used visual representations of model residuals to check that the assumptions of regression modelling were met in all cases. A single measurement of conjugated 11KT was excluded from all relevant analyses as it was far outside the range of all other values, and exerted large influence on the fitted model when included. Note that we control for any effects of sex in the statistical models and also include body mass as a covariate to account for the expected relationship between size and hormone release rates. We do not adjust observed endocrine state for size prior to modelling since this causes problems of interpretation for repeatability estimates (Wilson, 2018). Data handling and visualisation in R was carried out using the ‘tidyverse’ packages (Wickham, 2017). R code for performing the analyses is provided in the supplementary materials.

#### Habituation effects on mean hormone levels

For each of the endocrine response traits (Cort_free_, 11KT_free_, Cort_conjugate_, 11KT_conjugate_) in turn, we tested for the hypothesised effects of habituation on hormone levels across repeated assays by fitting assay repeat (as a categorical variable with levels 1 to 4). We also included sex, and the interaction of sex and assay repeat to account for any sex difference in mean endocrine state or in average habituation. We specified assay repeat as a factor rather than as a continuous predictor to allow maximum flexibility (i.e., to avoid imposing a linear response in the mean). Tank (2-level categorical variable), sampling order (the order transferred from the holding tank to the individual beakers, fitted as a mean-centred linear effect), and body mass were included as further covariates to control for potential sources of variance not relevant to our present hypotheses. We standardised body mass measurements by centring and scaling (subtracting the population mean and dividing by the standard deviation) to improve their interpretability (Gelman and Hill, 2007; Schielzeth, 2010). We used conditional Wald F-tests to test all fixed effects, and reduced the model by dropping the sex × assay repeat interaction if not significant. For fixed effects inference we use a ‘random slopes’ model (described below) where possible, which groups repeated measures to avoid pseudoreplication and also allows individuals to vary as a continuous function of stressor exposure, preventing inflation of Type I and Type II errors when estimating population-level effects (Schielzeth and Forstmeier, 2009). In cases where the random slopes model did not converge, we used a random intercepts model instead.

#### Among-individual variation

First, for each response trait in turn we tested for among-individual variance within the reaction norm framework. For each response trait we use the simplified fixed effects structure as determined by the model used to test for habituation described above. To test for repeatable differences in average hormone levels (i.e., among-individual variance in reaction norm intercept) across all four repeats, we compared a ‘null’ model (with no random effects) to one in which we fit a random effect of individual ID. We then extended the random intercepts model to include among-individual differences in habituation rate (i.e., reaction norm slopes), by fitting both a random ID effect and a random interaction of ID with assay repeat number (this time as a continuous covariate), as well as the intercept-slope covariance. We compared nested models using likelihood ratio tests (LRTs), in which we assume that twice the difference in model log-likelihoods conforms to a chi-square distribution where the degrees of freedom are set by the number of additional parameters in the more complex model. When testing the effect of a single variance term, the test statistic distribution is assumed to correspond to a 50:50 mix of chi-squared distributions having 0 and 1 degrees of freedom respectively (Self and Liang, 1987; Visscher, 2006).

Second, for each response variable we formulated a multivariate (4-’trait’) model to test hypotheses about variance in-and covariance among-the four repeat-specific observations. Rather than using the raw data, we estimated (co)variances conditional on the repeat-specific means as well as fixed effects as described above. We fitted a series of nested models to test hypotheses about the structure of individual variation. The first model estimated no covariances, and constrained the repeat-specific variances to be equal. Model 2 allowed the repeat-specific variances to differ, and model 3 extended model 2 by also estimating all covariances. We compare nested models using LRTs as detailed above. Model 2 vs model 1 therefore tests whether phenotypic variance (conditional on fixed effects) changes significantly across repeats (i.e., that the amount of variation among individuals in hormone state changes over the repeated exposures, suggesting variation in habituation), and model 3 vs model 2 tests for the existence of significant within-individual covariance structure (i.e., that some degree of repeatability exists). Model 3 estimates the within-individual covariance-correlation matrix (conditional on fixed effects), which we used as the basis for a parametric bootstrap method (described in Boulton *et al*. 2014; Houslay *et al*. 2017) to generate approximate 95% confidence intervals on all parameters. Inspection of these parameters and their associated 95% CIs can also be used to diagnose variation in habituation (again, evidenced by changes in observed endocrine state variance over sampling repeats, and/or declining (within-individual) correlation with increasing time between observations (such that *r*_1_,_2_>*r*_1,3_>*r*_1,4_)).

## Results

### Changes in hormone concentrations following repeated stress exposures

Both free cortisol and free 11KT show significant changes in their mean waterborne concentrations (i.e., immediately following transfer-related stress exposure) across the four successive stress exposures. Free cortisol concentration follows a pattern that largely conforms to our expectations of habituation, the mean declining significantly from stress exposures 1 to 2 and remaining stable at 3, although then showing a slight increase at exposure 4 (Fig. 2A, Table 1A). Females produce higher levels of cortisol than do males across all repeats. We find no sex differences in habituation rate for free cortisol (sex × stressor number interaction: F_3, 63.3_ = 0.63, *P* = 0.60). In contrast, male and female guppies do differ in how their 11KT levels change across repeated assays (sex × stressor number interaction: F_3, 65.9_ = 5.11, *P* = 0.003; Fig. 3, Table 1B). Males show fairly stable levels of 11KT across all repeats, while females show increased 11KT after the first assay. Males consistently produce greater levels of 11KT in comparison to females.

**Table 1:**
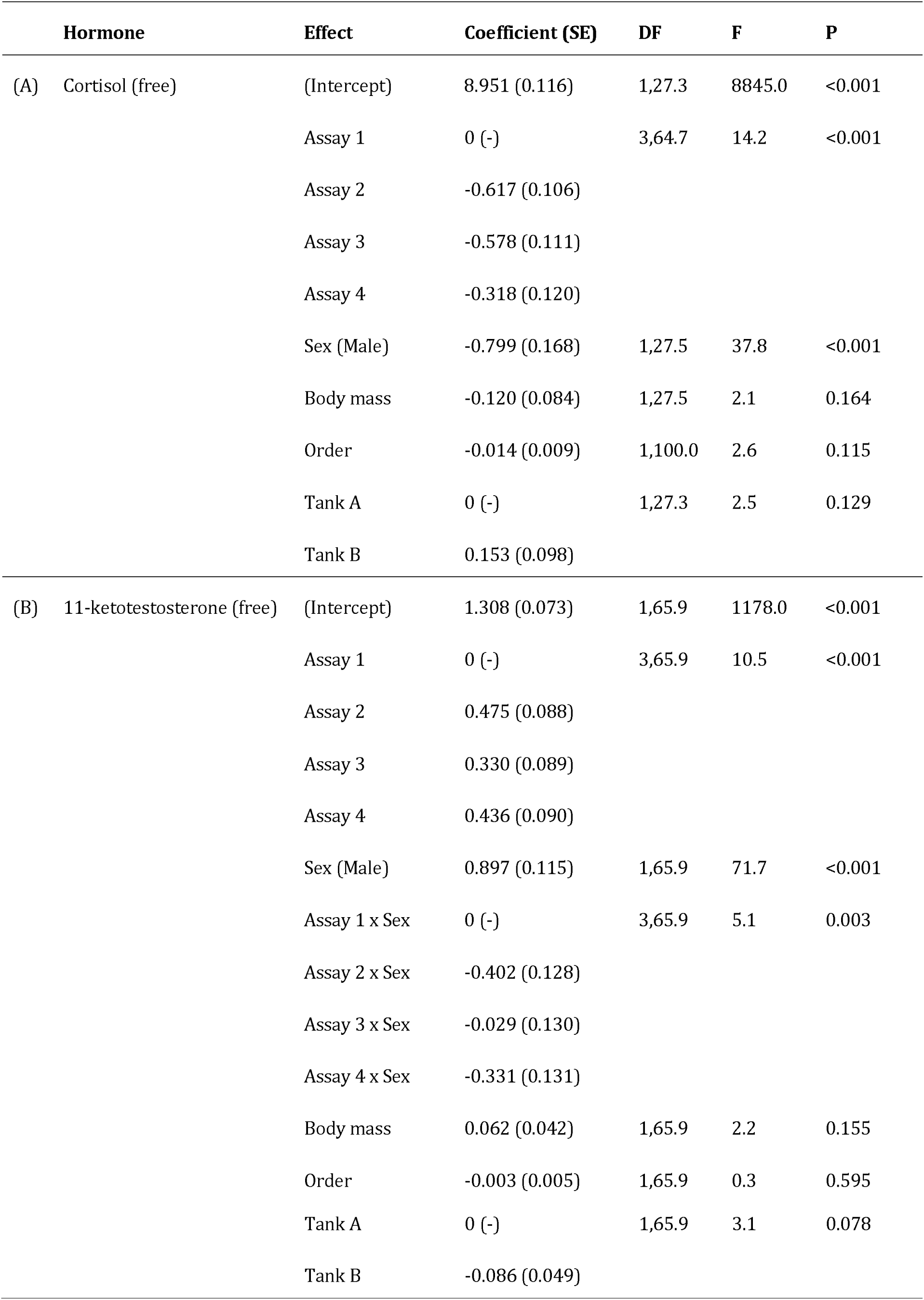
Fixed effect estimates from mixed-effects model analyses of free circulating (A) cortisol and (B) 11-ketotestosterone levels in individual guppies over four repeated measures.

|  | **Hormone** | **Effect** | **Coefficient (SE)** | **DF** | **F** | **P** |
| --- | --- | --- | --- | --- | --- | --- |
| (A) | Cortisol (free) | (Intercept) | 8.951 (0.116) | 1,27.3 | 8845.0 | <0.001 |
|  |  | Assay 1 | 0 (-) | 3,64.7 | 14.2 | <0.001 |
|  |  | Assay 2 | -0.617 (0.106) |  |  |  |
|  |  | Assay 3 | -0.578 (0.111) |  |  |  |
|  |  | Assay 4 | -0.318 (0.120) |  |  |  |
|  |  | Sex (Male) | -0.799 (0.168) | 1,27.5 | 37.8 | <0.001 |
|  |  | Body mass | -0.120 (0.084) | 1,27.5 | 2.1 | 0.164 |
|  |  | Order | -0.014 (0.009) | 1,100.0 | 2.6 | 0.115 |
|  |  | Tank A | 0 (-) | 1,27.3 | 2.5 | 0.129 |
|  |  | Tank B | 0.153 (0.098) |  |  |  |
| (B) | 11-ketotestosterone (free) | (Intercept) | 1.308 (0.073) | 1,65.9 | 1178.0 | <0.001 |
|  |  | Assay 1 | 0 (-) | 3,65.9 | 10.5 | <0.001 |
|  |  | Assay 2 | 0.475 (0.088) |  |  |  |
|  |  | Assay 3 | 0.330 (0.089) |  |  |  |
|  |  | Assay 4 | 0.436 (0.090) |  |  |  |
|  |  | Sex (Male) | 0.897 (0.115) | 1,65.9 | 71.7 | <0.001 |
|  |  | Assay 1 x Sex | 0 (-) | 3,65.9 | 5.1 | 0.003 |
|  |  | Assay 2 x Sex | -0.402 (0.128) |  |  |  |
|  |  | Assay 3 x Sex | -0.029 (0.130) |  |  |  |
|  |  | Assay 4 x Sex | -0.331 (0.131) |  |  |  |
|  |  | Body mass | 0.062 (0.042) | 1,65.9 | 2.2 | 0.155 |
|  |  | Order | -0.003 (0.005) | 1,65.9 | 0.3 | 0.595 |

|  |  |  |  |  |
| --- | --- | --- | --- | --- |
| Tank A | 0 (-) | 1,65.9 | 3.1 | 0.078 |
| Tank B | -0.086 (0.049) |  |  |  |

**Figure 2:**
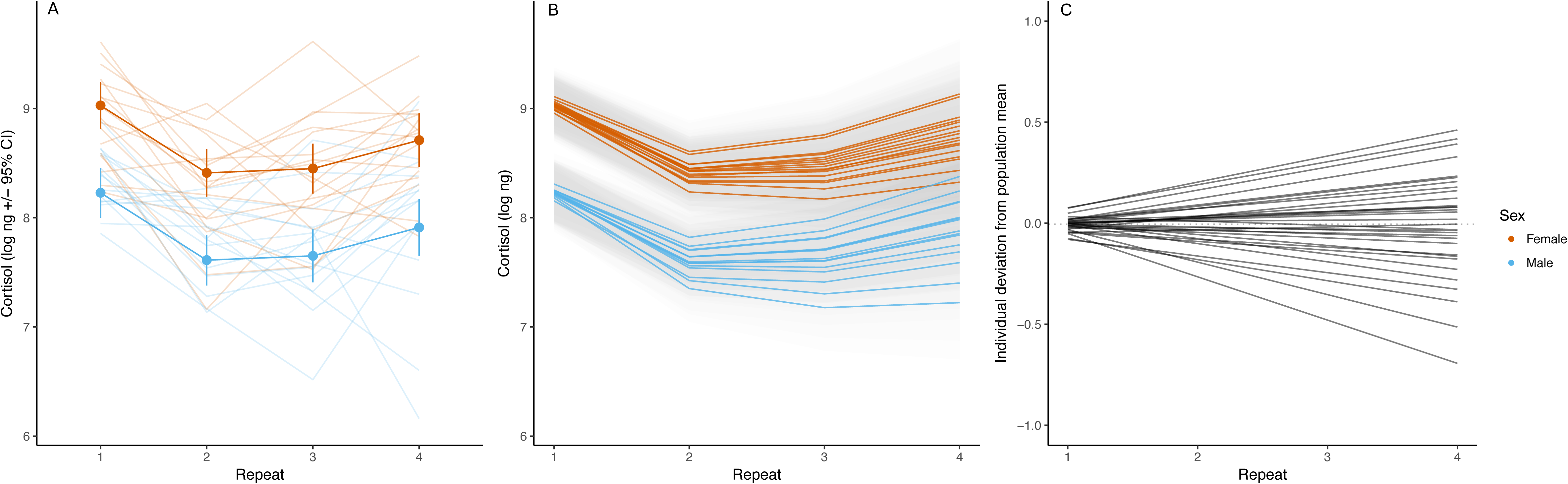
Variation in free circulating cortisol (log-transformed nanograms) in guppies. Panel (A) shows changes in cortisol as a function of stressor exposure (here, sampling repeat) separately for each sex; points are predictions from a linear mixed effects model (with 95% confidence intervals), with raw individual-level data in faint lines. Panel (B) shows predictions for each individual from a random regression model (coloured by sex; shaded area gives 95% confidence interval around each prediction), and (C) shows individual deviations from the population mean after accounting for all fixed effects (including sex and repeat).

**Figure 3:**
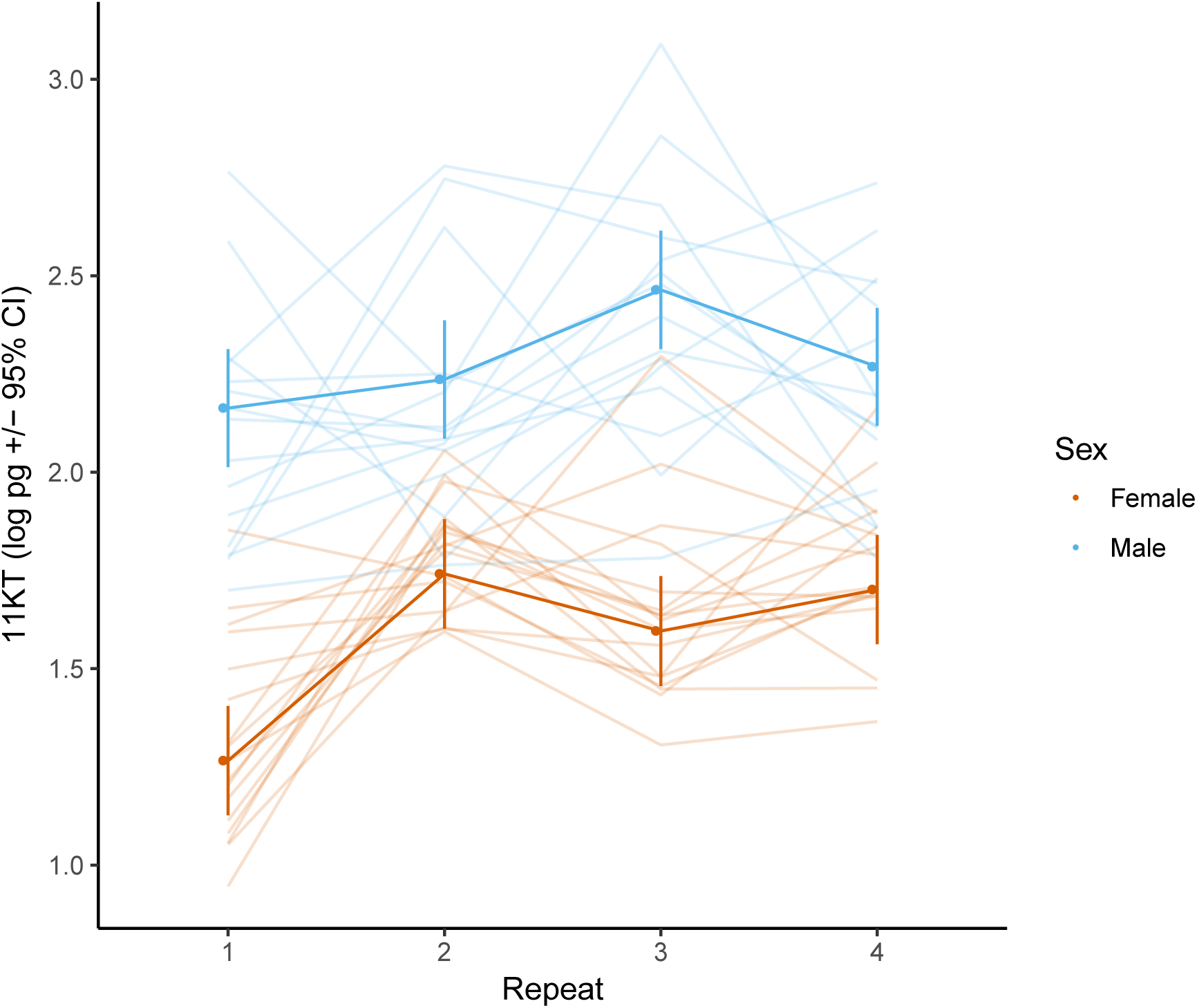
Variation in free circulating 11-ketotestosterone (log-transformed picograms) in guppies. Points show predictions from a linear mixed effects model (with 95% confidence intervals) as a function of stressor exposure (sampling repeat), plotted separately for each sex. Raw individual-level data is shown in faint lines.

### Reaction norm analyses of among-individual variation

LRT comparison of the univariate mixed models with and without a random effect of individual identity) provides statistical support for repeatable differences in average free cortisol concentrations across the repeated stress exposures. That said, the adjusted repeatability (i.e., the ratio of among-individual variation to phenotypic variation conditional on fixed effects) for free cortisol is relatively low (R=0.15 SE 0.10, χ^2^_0,1_=3.11, *P* = 0.04). Individuals do not show significant differences in their average level of free 11KT (R=0.03 SE 0.08, χ ^2^_0,1_ = 0.14, *P* = 0.36).

Addition of random reaction norm slope terms to the mixed models does not lead to a significantly better model for either free cortisol or free 11KT. For free cortisol, visual inspection of model predictions is certainly suggestive of an improved fit to the observed data (Figs 2B,C), though the apparent improvement is not significant (χ ^2^_2_ = 4.12, *P* = 0.13). Under this random slope model, the correlation between individual intercepts and slopes is strongly positive (such that individuals with higher average levels of cortisol also had a more positive slope), although the standard errors around the estimate are large (r = 0.88 SE 0.56). For free 11KT we were unable to detect any estimable variance in random slope (χ ^2^_2_ = 0, *P* = 1).

### Character state analyses of among-individual variation

For free cortisol, there is a significant change in repeat-specific variances, such that variance increases over successive repeats (model 2 vs model 1: χ ^2^_3_ = 8.2, *P* = 0.03; Fig. 4). This increase in variance over stress exposures suggests individuals may differ in their patterns of habituation. We also found significant within-individual covariance structure between repeats (model 3 vs model 2: χ ^2^_6_ = 14.0, *P* = 0.03), indicative of some repeatable variation among individuals. However, examination of the covariance-correlation matrix (Table 2A) shows that the correlations, though largely positive (5 out of 6) as predicted, are relatively weak (ranging from −0.239 to 0.345). Furthermore, there is no clear pattern of decline in the strength of the correlations as the interval between pairs of exposures increases. For free 11KT, we found no significant changes in repeat-specific variances for (model 2 vs model 1: χ ^2^_3_ = 0.6, *P* = 0.89), and no evidence for significant within-individual covariance structure (model 3 vs model 2: χ ^2^_6_ = 3.1, *P* = 0.62; correlations ranging from −0.184 to 0.197, Table 2B).

**Figure 4:**
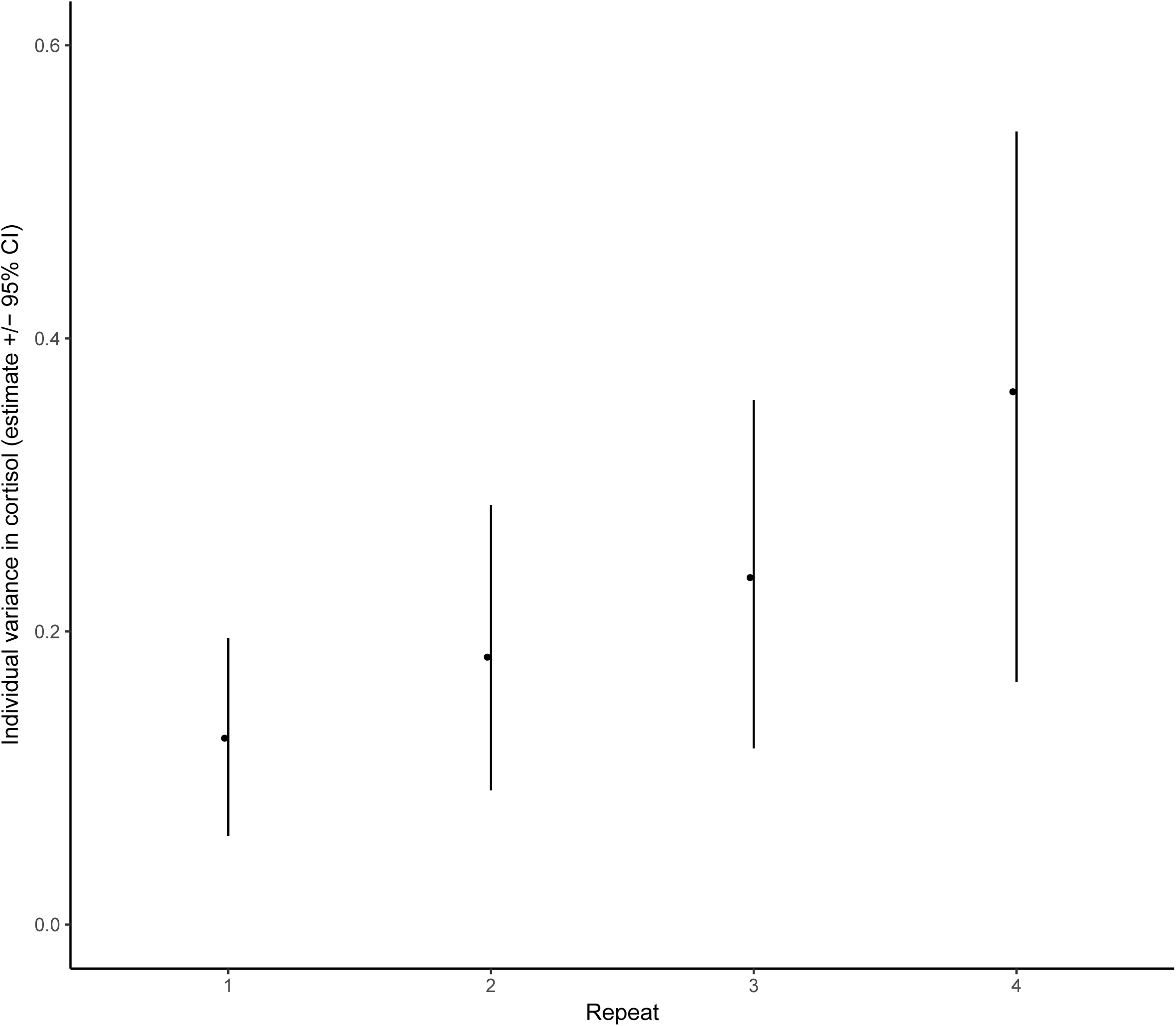
Change in individual variation in cortisol over increasing stressor exposures. Variance estimate and 95% confidence intervals (from 5000 bootstrapped replicates) for each sampling repeat, calculated from a multivariate mixed effects model fit with ASreml-R.

**Table 2:** Covariance-correlation matrix (conditional on fixed effects) for free circulating (A) cortisol, (B) 11KT. Variances are on shaded diagonals, covariances below and correlations above. In parentheses are the 95% confidence intervals on calculated from 5000 bootstrapped replicates.

|  |  |  |  |  |
| --- | --- | --- | --- | --- |
| A) Free cortisol | **Repeat 1** | **Repeat 2** | **Repeat 3** | **Repeat 4** |
| **Repeat 1** | 0.127 (0.058,0.189) | 0.121 (-0.505,0.327) | 0.345 (-0.074,0.486) | 0.141 (-0.45,0.352) |
| **Repeat 2** | 0.018 (-0.036,0.075) | 0.182 (0.087,0.28) | 0.34 (-0.099,0.479) | -0.239 (-1.262,0.077) |
| **Repeat 3** | 0.06 (-0.006,0.126) | 0.071 (-0.01,0.151) | 0.237 (0.114,0.353) | 0.265 (-0.217,0.429) |
| **Repeat 4** | 0.03 (-0.046,0.113) | -0.062 (-0.159,0.03) | 0.078 (-0.031,0.189) | 0.364 (0.183,0.549) |
| B) Free 11KT | **Repeat 1** | **Repeat 2** | **Repeat 3** | **Repeat 4** |
| **Repeat 1** | 0.076 (0.036,0.114) | -0.184 (-1.214,0.113) | -0.017 (-0.79,0.251) | 0.197 (-0.382,0.367) |
| **Repeat 2** | -0.013 (-0.039,0.012) | 0.063 (0.03,0.096) | 0.143 (-0.463,0.338) | -0.048 (-0.781,0.205) |
| **Repeat 3** | -0.001 (-0.028,0.028) | 0.01 (-0.015,0.035) | 0.076 (0.036,0.112) | 0.103 (-0.551,0.313) |
| **Repeat 4** | 0.013 (-0.013,0.037) | -0.003 (-0.023,0.019) | 0.007 (-0.018,0.031) | 0.059 (0.03,0.09) |

## Discussion

Our results show a striking pattern of change in the mean stress-induced free cortisol concentrations expressed by guppies across repeated stress exposures. This pattern was consistent with our predictions regarding habituation to the sampling protocol stressor following repeated exposure to it, with fish showing significantly higher free cortisol concentrations during the first exposure than during the following three. Free 11KT concentrations also change on average across the repeated stress exposures, most clearly in females which produce relatively low 11KT levels on the first (nominally more stressful) exposure relative to later ones. Our analyses of among-individual variation-conducted within both random regression and multivariate “character state” frameworks − also provide clear evidence that individuals differ in their average stress-induced hormone levels across sampling events. While we find some limited support for our final prediction that individuals would also differ in their rates of habituation with regard to stress-induced free cortisol release, statistical evidence is equivocal (potentially due to limited statistical power).

### Habituation in stress-induced free hormone concentrations

The high levels of (average) free cortisol during the first stress exposure, coupled with the decreased levels during subsequent exposures, are consistent with our prediction that habituation should occur with repeated stressor exposure. The population-level habituation dynamics seen here echo largely those reported previously in guppies (Fischer et al., 2014) and other small fish (e.g., Wong *et al*. 2008; Fürtbauer *et al*. 2015). This pattern constitutes a form of plasticity in stress responsiveness that is widely considered adaptive in the context of ‘coping’ with environmental challenges: while the stress response enables individuals to react appropriately to acute environmental change, habituation may be important to protect against the wear and tear that would arise from continued activation of the HPA/I axis. Less clear, however, is the reason for the apparent increase in the stress-induced free cortisol concentration levels between the third and fourth stress exposures (Fig. 2A). While the average level of free cortisol is still lower during the fourth stress exposure than the first-indicating that individuals are still habituated to repeated stress exposure-there is a small increase in the final stress exposure compared to the second and third. This pattern is suggestive of a more complex change in HPA/I axis performance in response to repeated stress exposure than initially envisaged, and could conceivably reflect a number of different physiological changes acting in isolation or concert. For example, while this pattern could reflect an increase in the magnitude of stress-induced cortisol release between exposures 3 and 4 (consistent with some form of attenuation of habituation), it could also arise in part from impacts of habituation on baseline cortisol concentrations prior to stress exposure (if baseline cortisol concentrations rose over the course of habituation, this too could contribute to an increase in the stress-induced cortisol concentrations). Teasing apart the relative contributions of these proximate explanations for the observed pattern would be fruitful prior to speculation on ultimate explanations.

We also find evidence of clear sexual dimorphism in levels of stress-induced free cortisol, with females producing more on average across all repeats. This endocrine sex difference cannot be attributed simply to sex differences in body mass (female guppies being much larger on average than males, and also more variable), as a linear effect of body mass was not significant in our model of free cortisol. This sex difference in stress responsive cortisol levels could therefore reflect different selection pressures acting on the male and female stress response, possibly due to sex differences in life histories, rather than simple associations with body size (Ricklefs and Wikelski, 2002). Male guppies have higher free 11KT concentrations than females across all stress exposures, in line with the androgenic role of this hormone. However, while the cortisol habituation dynamics are remarkably similar between males and females, 11KT shows sex-specific trajectories across repeat stress exposures. Male levels remain stable (on average) while females tend to have a depressed initial response before stabilising at a higher level. While increased cortisol concentrations have been shown to depress circulating 11KT concentrations in other (male) teleost fish (e.g. male *Salmo trutta*, Pickering et al., 1987), we see no evidence of a link between the stress-induced concentrations of the two hormones in male guppies. 11KT is important for male mating behaviour in teleosts generally (Borg, 1994); given the well-characterised sexual selection through female choice on male guppies (Brooks and Endler, 2001; Head and Brooks, 2006; Luyten and Liley, 1991), it seems plausible that a lack of stress-sensitivity reflects strong selection on males to maintain mating behaviours even in the face of high environmental stress. In females, by contrast, the low 11KT concentrations during the first stress exposure relative to the following exposures could indeed be consistent with GC-related inhibition of circulating 11KT concentrations: the strongest cortisol response in the first stress exposure could be depressing 11KT levels to a greater extent than the weaker cortisol responses to the subsequent exposures.

### Individual variation in stress-induced hormone levels and habituation

In addition to the population-level evidence of habituation reported above, we also find evidence of consistent differences among individuals in their average stress-induced free cortisol concentrations (after controlling statistically for variation in body size). Whether there are differences among individuals in their rate of habituation to the stressor is less clear. Plotting fitted reaction norms shows that individuals seem to diverge over repeated stress exposures, yet allowing for the existence of individual differences in these habituation gradients did not yield a model of significantly better fit to the data than assuming that all individuals shared the same habituation gradient. Similarly, the character state approach shows that individual variance increases over successive repeats; however, the covariance structure is not consistently positive, suggesting that this is not a simple case of individuals habituating at different rates. That these patterns are suggestive of individual variation in habituation, but without yielding conclusive evidence for it, points to a lack of sufficient power in our study. Mixed effects models-particularly those with complex random effects structures − can be data hungry, and it appears likely that a greater number of individuals would need to be assayed in order to demonstrate that there are significant differences among individuals in their rate of to habituation to a repeated stressor (Martin et al., 2011). Similar (but higher-powered) studies are required to shed light upon the causes and consequences of individual variation in endocrine traits.

One of the main drivers behind the interest in among-individual variation in labile traits is that its presence means there is an opportunity for selection to act on them. Such variation can stem from a variety of different sources, which have been discussed widely in the behavioural literature (e.g., Bierbach et al., 2017; Fisher et al., 2018; Reale et al., 2010) and are liable to apply to the endocrine stress response as well. For example, previous studies in guppies alone have shown effects on the GC response due to developmental plasticity (Chouinard-Thuly et al., 2018) and environmental stressors (Fischer et al., 2014). Recent studies of the population used here have shown that stress-related behaviours also respond plastically to environmental change (perceived level of predation risk) but show limited among-individual variation in this plasticity (Houslay et al., 2018). We also know that that these behaviours are under some degree of genetic control (White and Wilson, 2018; White et al., 2018), which may or may not be the case for cortisol expression. In fact, some debate exists as to whether hormone levels themselves can be viewed as heritable ‘traits’ under selection (Hau and Goymann, 2015; Zera et al., 2007), but there is certainly increasing evidence of a genetic basis to GC variation (Jenkins et al., 2014; Stedman et al., 2017). Links between GC baseline and/or reactivity and individual fitness have also been documented in a number of species (Blas et al., 2007; Cabezas et al., 2007; Patterson et al., 2014; Vitousek et al., 2014; Vitousek et al., 2018). Williams (2008) advocated looking beyond the ‘tyranny of the Golden Mean’ to embrace individual variation in endocrine systems; this challenge is starting to be taken up, with exciting possibilities for our understanding of the evolution of flexible phenotypes and hormonally mediated suites of traits (discussed further in Cockrem, 2013; Hau et al., 2016; Hau and Goymann, 2015; Taff and Vitousek, 2016).

### Conclusions

As predicted, we find that guppies habituate quickly to repeated exposure to a stressor. We also provide examples of a framework for characterising GC variability and flexibility within and among individuals using both reaction norm and character state approaches. Our results show individual variation in stress-induced cortisol levels, in addition to some limited evidence of individual variation in habituation to the stressor. Such variation provides material for selection to act upon, and could (given underlying genetic variation in habituation rate) enable a population to mount an evolutionary response-for example, to the deleterious effects of a failure to habituate to chronic stress. Given the current lack of consensus around how to diagnose chronic stress in populations (Dickens and Romero, 2013), one fruitful line of future research could be to estimate covariance at the among-individual or genetic level between failure to habituate to repeated exposure to a predictable, homotypic stressor (such as the sampling process itself) and a loss of body mass, changes in behaviour, or even survival. More broadly, our study adds to the burgeoning body of research showing that individuals differ in flexible endocrine traits that are likely to influence fitness. Future challenges include the investigation of whether individual habituation rates are themselves repeatable, and whether genetic links exist between GC variation and other aspects of the phenotype on which selection acts.

## Acknowledgements

We are grateful to A.J. Grimmer for assistance with animal husbandry, and to P. Sharman for help in the lab.

## Data statement

The data used in this study will be uploaded to Dryad upon completion. R code for analyses is provided in the supplementary materials.

## Appendix A: Analysis of conjugated fractions of cortisol and 11-ketotestosterone

### Mean effects

We find significant effects of repeated stress exposure on conjugated hormone fractions. For the conjugated form of cortisol this appears to be due largely to increased concentrations at exposures 3 and 4 relative to earlier exposures (Table A1a). We find no sex differences in how these levels changed over the repeated measures (sex × stressor number interaction: F_3,62.9_ = 0.91, *P* = 0.44). We also find no clear sex differences in mean conjugated cortisol concentrations, although there is a positive effect of mass. Concentrations of the conjugated form of 11KT also change over repeats, rising to a peak at the third exposure before decreasing in the final exposure (Table A1b). Males exhibit consistently higher levels of conjugated 11KT, and there are no sex differences in the response to repeated assays (F_3,88.7_ = 0.86, *P* = 0.46).

### Among-individual variance: random regression

Under the random regression approach, there is significant variance among individuals in the conjugated forms of both hormones. We find moderate adjusted repeatability for conjugated cortisol (R = 0.26 SE 0.10, χ ^2^_0,1_ = 8.91, *P* = 0.001) and for 11KT (R = 0.27 (SE 0.10), χ ^2^_0,1_ = 9.30, *P* = 0.001). For conjugated cortisol the comparison of random intercept and random slope models is non-significant (χ ^2^_2_ = 1.64, *P* = 0.44). For conjugated 11KT we do not detect any estimable variance in random slope (χ ^2^_2_ = 0, *P* = 1).

### Among-individual variance: character state

Using the character state approach, conjugated cortisol shows no significant change in variance over repeats, but there is significant within-individual covariance structure. The covariance-correlation matrix shows that correlations between pairs of assays are weakly positive (ranging from 0.131 to 0.411, Table A2a), and we also find a qualitative pattern of declining strength as the inter-observation time increases. Conjugated 11KT does show changes in variance across repeats (χ ^2^_3_ = 33.3, *P* < 0.001), as well as within-individual covariance structure (χ ^2^_6_ = 19.8, *P* = 0.003) and a pattern of positive correlations among repeats (ranging from 0.124 to 0.561, Table A2b), which tend to decline in strength with increasing inter-observation interval.

**Table A1:**
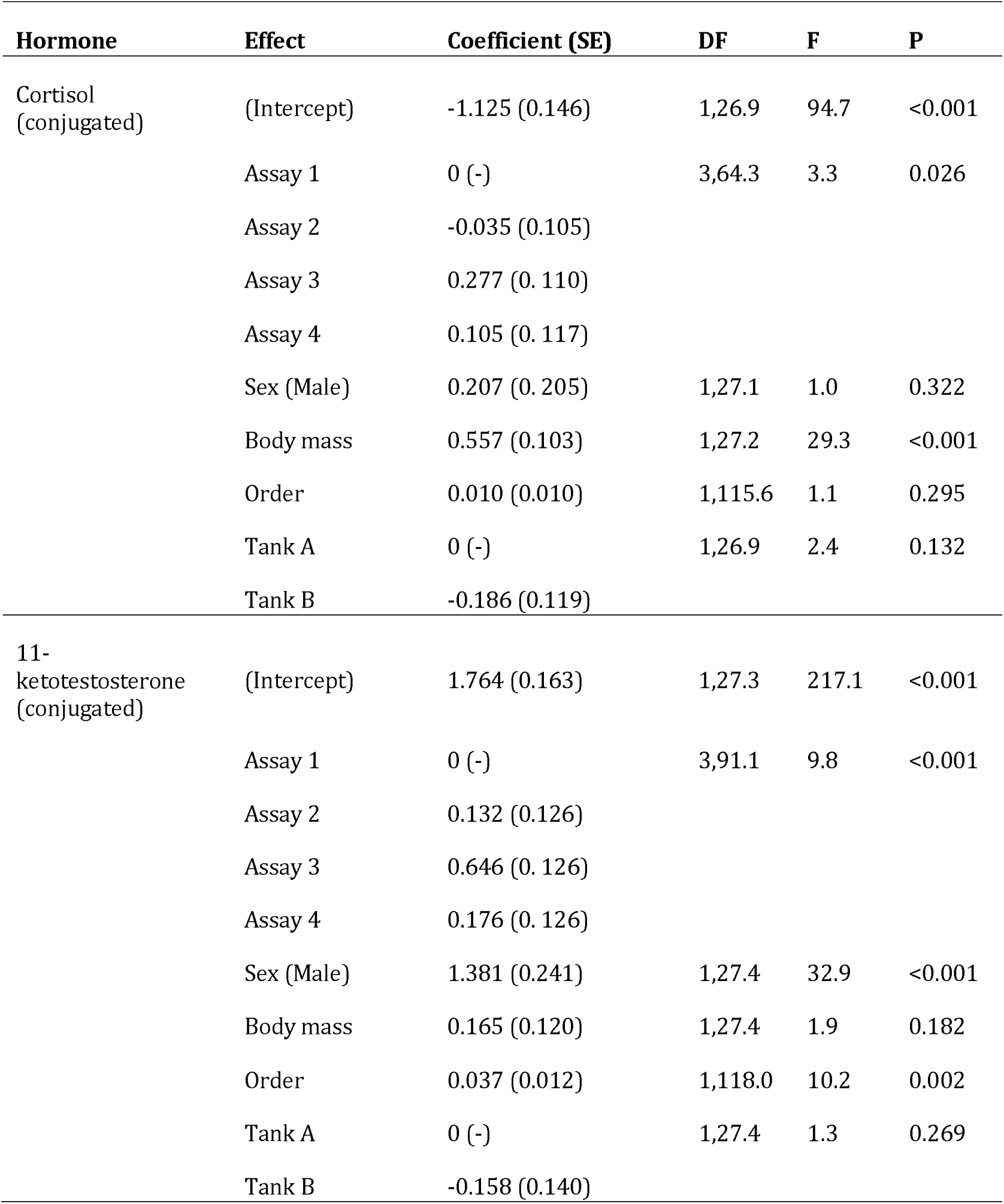
Fixed effect estimates from mixed-effects model analyses of conjugated (a) cortisol and (b) **11**-ketotestosterone levels in individual guppies over four repeated measures.

**Table A2:** Covariance-correlation matrix (conditional on fixed effects) for conjugated circulating (a) cortisol, (b) **11KT**. Variances are on shaded diagonals, covariances below and correlations above. In parentheses are the 95% confidence intervals on calculated from 5000 bootstrapped replicates.

| c) Conjugated cortisol | Repeat 1 | Repeat 2 | Repeat 3 | Repeat 4 |
| --- | --- | --- | --- | --- |
| Repeat 1 | 0.238 (0.116,0.367) | 0.411 (-0.008,0.519) | 0.202 (-0.394,0.356) | 0.131 (-0.499,0.327) |
| Repeat 2 | 0.126 (-0.001,0.245) | 0.396 (0.198,0.609) | 0.379 (-0.04,0.492) | 0.162 (-0.376,0.368) |
| Repeat 3 | 0.045 (-0.042,0.122) | 0.109 (-0.005,0.217) | 0.208 (0.097,0.321) | 0.367 (-0.052,0.508) |
| Repeat 4 | 0.029 (-0.053,0.11) | 0.046 (-0.052,0.16) | 0.075 (-0.005,0.16) | 0.201 (0.098,0.309) |

| d) Conjugated 11KT | Repeat 1 | Repeat 2 | Repeat 3 | Repeat 4 |
| --- | --- | --- | --- | --- |
| Repeat 1 | 0.231 (0.105,0.346) | 0.477 (0.113,0.582) | 0.176 (-0.404,0.371) | 0.124 (-0.6,0.31) |
| Repeat 2 | 0.157 (0.018,0.292) | 0.471 (0.246,0.729) | 0.267 (-0.184,0.439) | 0.372 (-0.034,0.517) |
| Repeat 3 | 0.1 (-0.105,0.313) | 0.215 (-0.074,0.537) | 1.381 (0.646,2.053) | 0.561 (0.273,0.645) |
| Repeat 4 | 0.029 (-0.062,0.109) | 0.123 (-0.005,0.264) | 0.319 (0.07,0.553) | 0.233 (0.102,0.357) |

